# Activation of mTOR by release of extracellular cholesterol stores controls the transition from quiescence to growth in *C. elegans*

**DOI:** 10.1101/2022.08.26.505407

**Authors:** Kathrin Schmeisser, Damla Kaptan, Bharath Kumar Raghuraman, Andrej Shevchenko, Sider Penkov, Teymuras V. Kurzchalia

## Abstract

Recovery from the quiescent developmental stage called dauer is an essential process in *C. elegans* and provides an excellent model to understand how metabolic transitions contribute to developmental plasticity. We here show that the depletion of sterol-binding proteins SCL-12 and SCL-13 is the key change in the *C. elegans* proteome in early dauer recovery. This process releases a cholesterol store that is sequestered in the gut lumen during the dauer state to facilitate the transition into reproductive development. First, the stored cholesterol undergoes endocytosis into the lysosomes of the intestinal cells, where it activates mTOR to promote protein synthesis and growth. Second, it is used for the production of dafachronic acids that switch metabolic programs at the transcriptional level. These processes are essential for population fitness and survival, as loss of SCL-12 and SCL-13, depletion of sterols, and loss of mTOR precludes quiescence exit, ultimately leading to the expiration of the entire population.

## Introduction

Environmental conditions such as temperature, population density, or the presence of other species, amongst many others, impact the development of an organism at different levels. The most potent exogenous stressor is the availability of nutrients and micronutrients, which has a dramatic effect on the growth of almost all species. However, the lack of essential nutrients and other environmental pressures are common and have led to multiple adaptation strategies amongst the species such as quiescence or alternative developmental programs. This so-called developmental plasticity ensures the survival of the organism during harsh times by adjusting growth and physiology until conditions improve [1].

*C. elegans* is an excellent model to study developmental plasticity as it adapted several strategies to react to environmental stress. The free-living nematode relies mostly on microorganisms in organic material as food source that they find in the soil, their natural habitat in the wild [2]. Their development is regulated by many conserved signaling pathways, resulting in a robust and invariant pattern of cell divisions and a steady cell number of 959 in all animals [3]. However, at times of averse environmental conditions, their development can slow or completely arrest in quiescent stages, as worms have several diapauses implemented in their life cycle: The L1 diapause [4] and the adult reproductive diapause [5], as well as the dauer state [6], which is an alternative developmental step: Late stage L1 larvae can alter their developmental trajectory and postpone reproduction by going through a preparatory second larval stage (L2d) and a quiescent dauer larval stage. This increases the chance of survival until environmental conditions improve and progeny can be sustained. Distinct dauer-inducing pheromones, which are short-chain ascarosides that the animals continuously release, facilitate dauer development. When levels of dauer-inducing pheromones in a population rise over a certain threshold, *C. elegans* will activate the dauer program [7]. Several signaling mechanisms, including the insulin/IGF-1, TGF-β, cyclic GMP, and steroid-signaling pathways control the ensuing significant changes in gene expression that lead to the characteristic morphological and metabolic modifications associated with this transition. The morphological changes include radial constriction of the cuticle, reduced body diameter, and a sealing up of the pharynx; all of which are adaptions to survive harsh environmental conditions. Their metabolism is substantially different than in reproductive larvae; they do not feed and rely on a catabolic mode of metabolism, using stored energy reserves. To save on these reserves, dauers are hypometabolic, which implies that heat production, aerobic respiration, and TCA cycle activity are significantly reduced [8]. At the same time, the glyoxylate shunt and gluconeogenesis are used for glucose generation from stored lipids [9, 10]. Dauers can survive in this state for weeks to months, surpassing their life expectancy in reproductive mode. Downstream of the insulin/IGF-1, TGF-β, cyclic GMP, and steroid- signaling pathways, two major transcription factors determine the developmental states on the gene expression level: the nuclear hormone receptor DAF-12 and the FoxO member DAF-16 [10]. The latter is mainly affected by insulin-like signaling via the insulin receptor orthologue DAF-2. DAF-12 is controlled by dafachronic acids (DA), which are bile acid-like steroid hormones that can exist in four isomers: two regioisomers, Δ4- and Δ7-DAs, whereas each of which exist as 25R- and 25S-diastereomers. The last, critical step in their synthesis is performed by DAF-9, a cytochrome P450 that oxygenates the side chain of cholesterol. In its DA-bound state, DAF-12 promotes reproductive development and suppresses dauer formation or facilitates dauer exit. When DA is not bound to DAF-12, dauer formation is induced [11, 12]. This, however, requires very low or even absent DA levels in the cellular environment, because DA acts in extremely low concentrations, probably in the picomolar range. It is still not fully understood how an absence of DA can be achieved, as cholesterol is ubiquitous in the environment and therefore also within the animal itself. The sterol methylase STRM-1 has been shown to reduce the amount of DA by modifying cholesterol in a manner such that DAF-9 cannot use it as substrate to generate DA. As a result, STRM-1-dependent methylation of sterols ensures that DA levels are maintained low, facilitating the regulation of dauer formation. However, only up to 50% of the sterol pool can be methylated [13]. As *C. elegans* cannot synthesize steroids *de novo* and need to obtain them from their diet, another way to lower DA levels could be the limiting of cholesterol digestion or compartmentalization/sequestration within the worm. On the other hand, such a compartmentalization mechanism has not been described yet, and given the ubiquitousness of cholesterol in the environment, it is almost impossible to avoid. This is demonstrated by the rigorous laboratory protocol that is needed to remove sterols from the components that generally constitute *C. elegans* laboratory cultures [12]. Such experiments, however, have helped to understand more about the role of cholesterol for *C. elegans* besides regulation of the dauer diapause. If kept under completely sterol-free conditions, the first generation of worms can still grow from eggs to adulthood. In the second generation though, they develop into dauer-like larvae with incomplete molting (so-called L2*). Thus, cholesterol is necessary for development through the larval stages, growth, and the correct shedding of old cuticles during molting processes, but only once its stores are completely depleted [14, 15].

In mammalian cells, it has been shown that cholesterol drives mTOR Complex 1 (mTORC1) recruitment and activation at the lysosomes to facilitate growth in response to nutrient cues [16]. mTOR is a highly conserved serine/threonine protein kinase and part of the catalytic subunit of two protein complexes, mTORC1 and mTORC2 [17]. Whereas both complexes can sense environmental conditions and regulate different cellular processes, mTORC1 is responsible for protein synthesis and growth and therefore of particular interest in this study. The major components of mTORC1 are the TOR kinase and the Regulatory Associated Protein of mTOR (RAPTOR), with LET-363 and DAF-15 being the *C. elegans* orthologues [18]. It has been shown that both LET-363 and DAF-15 are required during development to progress through the larval stages [19]. Interestingly, knockout of *daf-15*/RAPTOR causes larval arrest at the L3 stage, which, in terms of developmental timing, coincides with the dauer diapause that is regulated by the steroid hormone pathway [20]. Hence, it is possible that there is a cross-talk between mTORC1 and the steroid hormone pathway based on their shared sensitivity to cholesterol levels and developmental timing factors. However, such a cross-talk has not yet been investigated. Here, we describe a novel mechanism for cholesterol mediating the transitions between the dauer state and reproductive development that is dependent on a sequestration and compartmentalization of cholesterol during the dauer stage, potentially to minimize DA production. When worms enter dauer, cholesterol is bound to two proteins, SCL-12 and SCL-13, and stored in the intestinal lumen during the entire duration of the dauer stage. Upon dauer exit, cholesterol is released, facilitating the switch into the reproductive state in two distinct ways. First, it serves as precursor of DA, binding DAF-12, which subsequently switches developmental programs on a transcriptional level. Second, some of the sequestered cholesterol is transported by SCL-12 and SCL-13 into the lysosomes, where it signals to mTORC1 to boost growth. Furthermore, we show that mTORC1 and cholesterol in *C. elegans* can exert their growth-promoting effects only when both of them are present. This is, to our knowledge, the first evidence of cholesterol-mTOR interaction to enhance growth in an animal model, and elucidates some of the remaining questions about how cholesterol modulates development in *C. elegans*.

## Results

### Degradation of SCL-12 and SCL-13 is a major proteomic change during the early phase of dauer exit

To investigate early proteomic changes during the transition between the quiescent dauer larva and the reproductive state, we performed a 2D-differential gel electrophoresis (2D DIGE). We compared the protein landscape of wild-type dauers obtained from overcrowded medium before (labelled red) and during exit from the dauer state induced by exposure to food at low population density (labelled green). The most striking difference was two abundant proteins, which completely disappeared during the first 4 hours of dauer exit (two red spots in fig. 1A; marked by arrows). These were SCL-12 and SCL-13, which had been identified before as dauer-specific [21]. To obtain a more accurate picture of the proteomic switch, we performed label-free LC-MS/MS quantification of proteins in a strain that harbors the *daf-2(e1370)* allele. These worms form dauers at a restrictive temperature of 25°C and exit from the dauer state when shifted to the permissive temperature of 15°C, providing an easy model to switch between metabolic states [22]. Compared to the initial dauer state, exiting dauers 24 h after the switch to 15°C displayed 142 significantly differentially expressed proteins (fig. 1B), of which 110 were up- and 32 downregulated, respectively. Among the most upregulated proteins were a chromatin remodeller (GFI-1), a ribosomal protein involved in translation (RPL-28), and S-Adenosyl methionine synthetase (SAMS-1). The proteins with the strongest decrease in concentration were SCL-12 and SCL-13 (fig. 1B).

**Figure 1:**
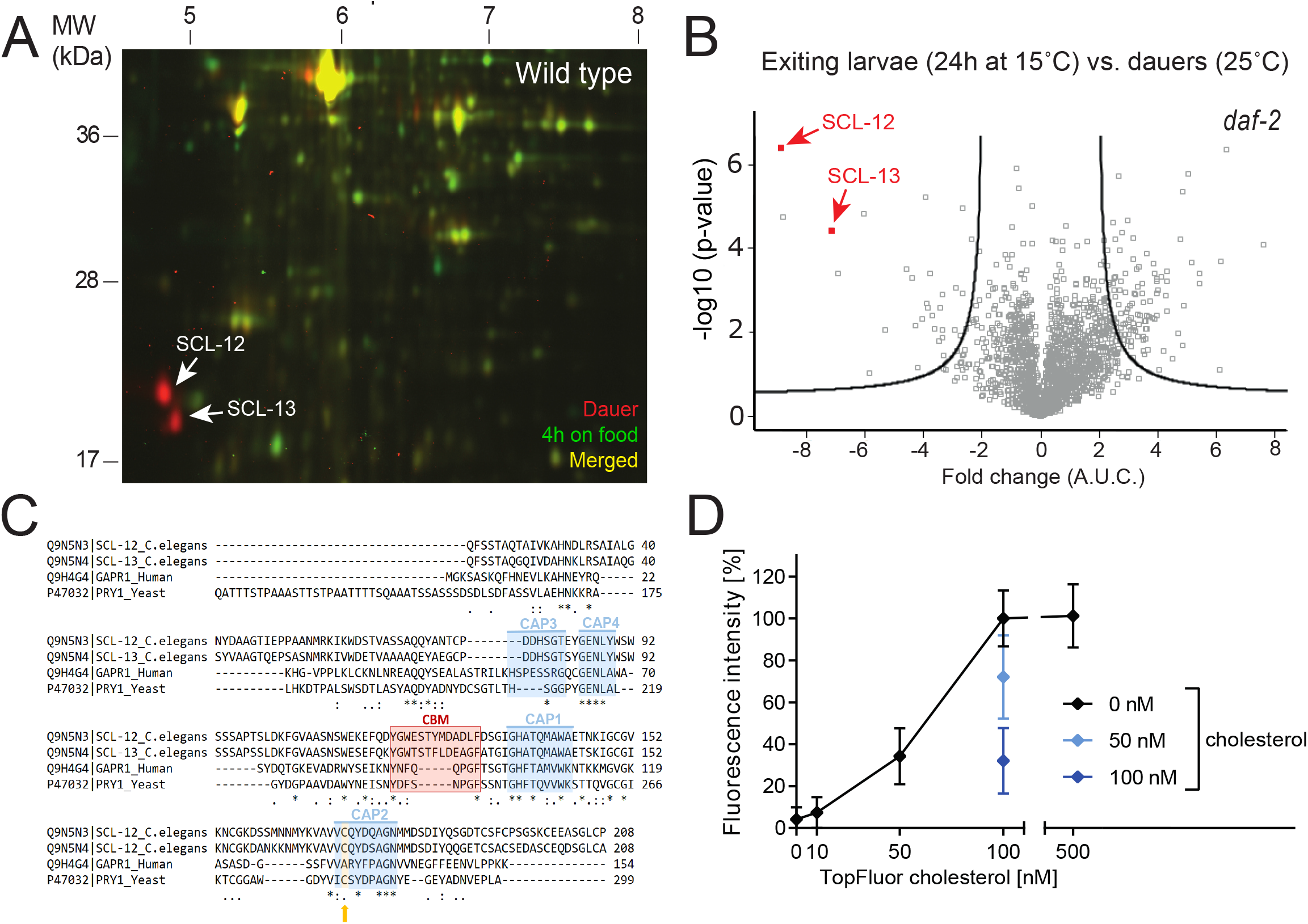
Degradation of SCL-12 and SCL-13 cholesterol-binding proteins is a major proteomic change during the early phase of dauer exit. **A** Overlay of false-colored 2D-DIGE images comparing the wildtype (wt) starvation dauer proteomes before (red) and 4 h after (green) introduction of food. Yellow spots represent proteins with no major changes in concentration. The most striking changes were SCL-12 and SCL-13 that are depleted within 4 h on food. **B** The proteome of 24 h exiting dauer *daf-2(e1370)* animals compared with the dauer proteome. Fold changes indicate upregulated (>0) and downregulated (<0) proteins in exiting dauer larvae. Black lines designate significance, i.e. p-values <0.05. **C** Alignment of *C. elegans* SCL-12 (UniProt ID: Q9N5N3_CAEEL), SCL-13 (Q9N5N4_CAEEL), human GLIPR2/GAPR1 (Q9H4G4) and yeast PRY1 (P47032) protein sequences. The functional signatures 1 to 4 of the CAP domain are highlighted in blue. Essential for cholesterol binding is the caveolin-binding motif (CBM; red) and a cysteine residue within the second signature of the CAP domain (orange). **D** Fluorescent intensity of purified SCL-12 protein that was incubated with various concentrations of TopFluor cholesterol (black) and pre-incubated with 50 nM (light blue) or 100 nM (blue) cholesterol for 1 h, bound to sepharose beads.

### SCL-12 and SCL-13 are conserved cholesterol-binding proteins

SCL(SCP-like)-12 and SCL-13 belong to a large family of proteins in *C. elegans* that display strong homology to the eukaryotic SCP/TAPS (Sperm-coating protein/Tpx/antigen 5/pathogenesis related-1/Sc7), or CAP (cysteine-rich secretory protein/antigen 5/pathogenesis related-1) superfamily. SCL-12 and SCL-13 align especially closely with the yeast CAP protein Pry1 (fig. 1C). Pry1 binds lipids, among them cholesterol [23, 24], and is used by yeast for the export of cholesterol acetate from the cytosol as part of a proposed lipid detoxification mechanism [25]. Other CAP proteins from arthropods and humans also bind cholesterol/lipids [26, 27]. The caveolin-binding motif (CBM) and a conserved cysteine residue of Pry1 are responsible for free cholesterol and cholesteryl acetate-binding [23, 24]. Within the CBM, the aromatic amino acids are essential to bind cholesterol [23]. SCL-12 and SCL-13 also possess a CBM with the respective aromatic residues, and a conserved cysteine in signature 2 of the CAP domain (C171; fig. 1C).

We tested whether *C. elegans* SCL-12 has sterol-binding properties by performing an *in vitro* binding assay [23]. Since SCL-12 and SCL-13 are highly homologous (76.8 % identity) and share the same promoter, we assumed that both proteins have nearly identical biochemical properties and thus, as representative for the *in vitro* binding assay, the sterol-binding of SCL-12 alone was interrogated. The protein was expressed in insect cells and after purification was incubated with a fluorescent analog of cholesterol (TopFluor cholesterol). As shown previously, this reagent behaves similarly to native cholesterol regarding intracellular transport or its incorporation into membranes [28]. The binding of TopFluor cholesterol to SCL-12 is concentration-dependent (fig. 1D). The specificity of binding to cholesterol was further supported using a competitive approach where SCL-12 was pre-incubated with unmodified cholesterol, which decreased the binding of TopFluor cholesterol significantly (fig. 1D). Thus, SCL-12 and SCL-13 are conserved sterol-binding proteins in *C. elegans* that bind cholesterol and perhaps other related sterols.

### SCL-12 accumulates in the dauer gut lumen and is transported back into the intestinal cells during dauer exit to be degraded

To understand where SCL-12 and SCL-13 exert their function, we generated a protein reporter in which SCL-12 is tagged with an mScarlet red fluorescent protein at its C-terminus and expressed it in the dauer-constitutive strain *daf-2* (SCL-12::mScarlet*;daf-2*; fig. S1A). Expression of the protein starts in the intestinal cells in late L2d worms (i.e. the specific L2 larval stage that precedes the dauer larva), about 44 h at 25°C after the embryo stage (fig. S1B). During dauer formation, fluorescence in the gut lumen increases (fig. S1C), and in fully formed dauers SCL-12 is exclusively found within the intestinal lumen (fig. 2A). We initially observed that SCL-12 and SCL-13 proteins are degraded during dauer exit (fig. 1A and 1B). Indeed, when exit was induced by temperature switch in SCL-12::mScarlet;*daf-2* animals, the gut lumen fluorescence decreased dramatically and almost disappeared within 24 h (fig. 2B; quantification of the process in fig. 2C). In parallel, faint fluorescence in the cytosol of the intestinal cells in punctate structures became visible (fig. 2B; 12 and 24 h time points). This fluorescence, however, also disappeared completely after 48 h. The same can be observed in SCL-12::mScarlet dauers with a wt background, where dauer exit was induced by introduction of food, although the process occurs much faster (fig. S1D, S1E).

**Figure 2:**
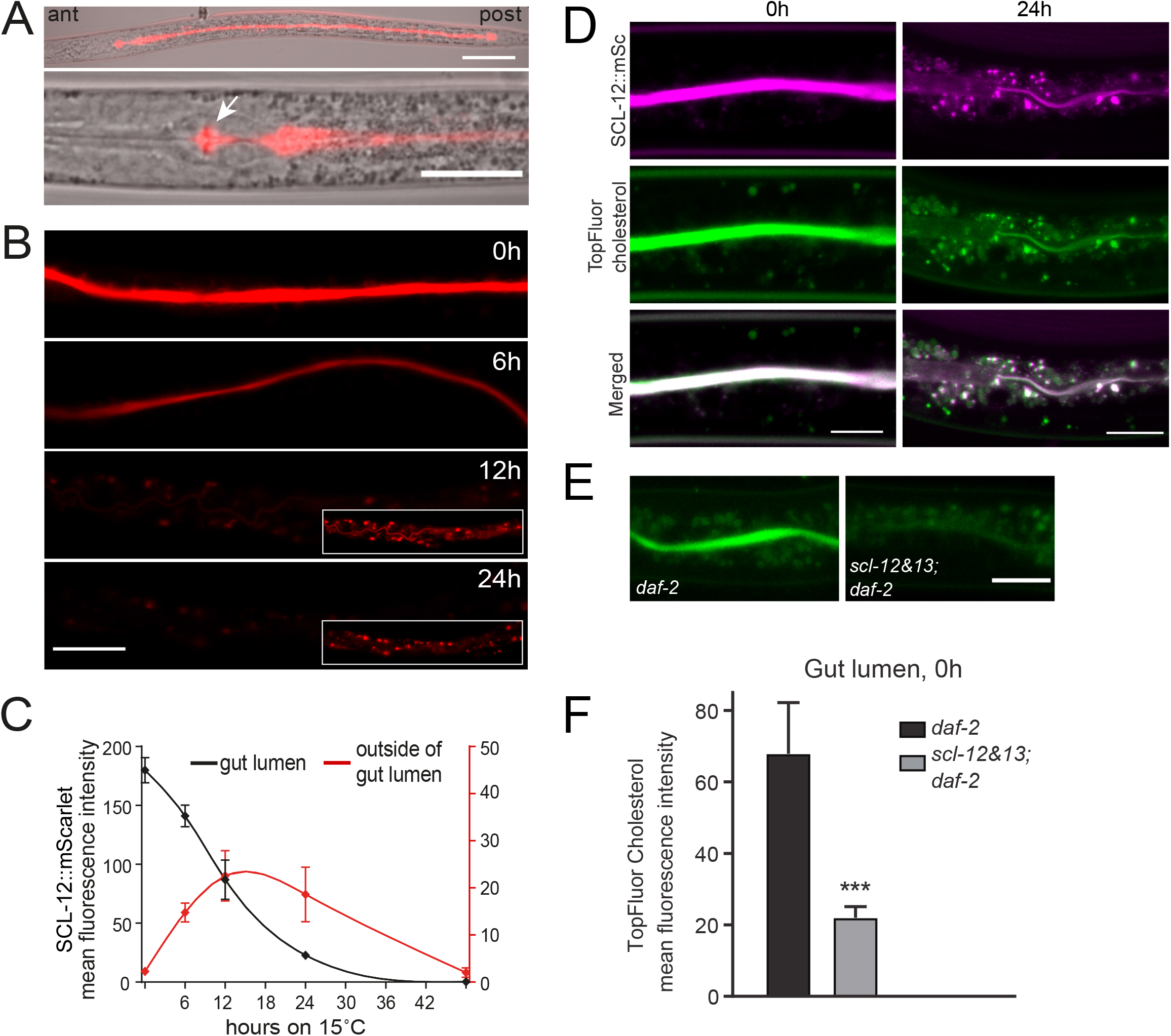
SCL-12 accumulates in the dauer gut during the dauer state where it sequesters cholesterol. **A** SCL-12::mScarlet reporter dauer larva, whole animal (top), head region (bottom), posterior (post) and anterior (ant) directions are marked. Scale bar top: 50 μm, bottom: 25 μm. **B** SCL-12::mScarlet;*daf-2* reporter dauer (0 h) and while exiting dauer at 6 h, 12 h and 24 h after temperature switch to 15°C. 12 h and 24 h insets are brightness adjusted images. Scale bar: 20 μm. **C** Mean fluorescence intensity of SCL-12::mScarlet;*daf-2* reporter dauers (0 h) and while exiting dauer after temperature switch to 15°C. Black: Fluorescence in the gut lumen. Red: Fluorescence outside of the lumen. **D** SCL-12::mScarlet reporter animals (SCL-12::mSc; magenta) and fluorescent TopFluor cholesterol (green) in the gut of a dauer larva (0 h; left panels) and 24 h after induction of dauer exit (right panels). Merged channels show a co-localization of both fluorophores. Scale bars: 10 μm. **E** *daf-2* and *scl-12&13;daf-2* dauers fed with TopFluor cholesterol prior entering the dauer state. Scale bar: 15 μm. **F** Mean fluorescence intensity of TopFluor cholesterol in the gut lumen from **E**, *daf-2* (black) and *scl-12&13;daf-2* (gray).

### SCL-12 and SCL-13 sequester cholesterol in the gut lumen of dauers

As SCL-12 and SCL-13 bind cholesterol *in vitro*, we wondered whether they exert an *in vivo* function in the transport and distribution of sterols as well. To visually follow the transport of cholesterol, we used the TopFluor cholesterol analog (the same compound that was used in the *in vitro* sterol-binding assay). SCL-12::mScarlet,*daf-2* worms were allowed to develop into dauer on plates, in which cholesterol was replaced by TopFluor cholesterol. Notably, like SCL-12, most of TopFluor cholesterol accumulates in the lumen of the dauer gut (fig. 2D, left panel). Moreover, it follows the SCL-12::mScarlet fluorescence during dauer exit into punctate structures and becomes less visible in the gut lumen (fig. 2D, right panel). This suggests that SCL-12 indeed binds cholesterol as worms enter the dauer state, sequesters it in the gut lumen during dauer, and releases it when degraded during dauer exit.

In order to clarify the role of SCL-12 and SCL-13 in the sequestration of cholesterol, we created a double-knockout strain introducing a 2.700 bp-deletion, spanning both the *scl-12* and *scl-13* genes on chromosome V (fig. S1A). *scl-12&13* mutants were crossed with *daf-2* animals, hereafter referred to as *scl-12&13;daf-2*. Like *daf-2*, this strain produces 100% dauers at the restrictive temperature (25°C), which, although being slightly smaller in size than *daf-2* alone (fig. S2A), do not display any visible phenotype. *scl-12&13;daf-2* mutants survive 1% SDS treatment, have the same oxygen consumption rate (OCR; fig. S2B; 0 h time point) and survival (fig. S2C). However, the distribution of TopFluor cholesterol in *scl-12&13;daf-2* dauers significantly differs from that of *daf-2* control animals and the gut lumen fluorescence is much lower (fig. 2E; quantification in 2F). Thus, we conclude that SCL-12 and SCL-13 are pivotal for the sequestration of cholesterol in the gut lumen of dauer larvae.

### Sequestration of cholesterol is needed for appropriate dauer exit

During maintenance of *scl-12&13*, we observed that dauer formation due to starvation is somewhat impaired. Yet, impaired dauer entry could not be observed in *scl-12&13;daf-2* mutants, most likely because *daf-2* at the restrictive temperature of 25°C prompts a dominant phenotype. However, when dauer exit is induced by temperature switch to the permissive 15°C in *scl-12&13;daf-2*, the mutants only partially recover, with about 40% of all animals remaining arrested (fig. 3A). Even in worms that do eventually exit dauer, growth is significantly delayed (fig. 3B) and they reach fertility two days later than the control (fig. S2D). Dauer exit depends on nutrient influx [29], and we asked whether delays in mouth opening/pumping could be responsible for the dauer exit phenotypes in *scl-12&13;daf-2*. However, no differences in in the onset of pumping compared to the control could be observed (fig. S2E), and only a slight decrease in the maximum pumping rate was detected 24 h after the switch to permissive temperature (fig. S2F). We therefore concluded that the arrest during/delay of dauer exit is independent of any defect in energy uptake. Moreover, a normal increase of OCR as worms exit dauer was observed in both *scl-12&13;daf-2* and *daf-2* mutants (fig. S2B), suggesting that the switch from dauer to reproductive metabolism/oxidative phosphorylation during dauer exit is unperturbed. We next aimed to understand whether cholesterol is a crucial molecule for dauer entry and exit. Therefore, we grew *daf-2* animals at 25°C on either control conditions (cholesterol^+^) or strictly sterol-free (cholesterol^-^), as previously described [12], and let them develop into dauer. We then induced dauer exit on cholesterol^+^ or cholesterol^-^ and scored recovered worms after 48 h (scheme in fig. 3C). As expected, worms recover normally when kept at cholesterol^+^ conditions at all times (cholesterol^+^/cholesterol^+^) and display impaired recovery when under cholesterol^-^ conditions (cholesterol^-^/cholesterol^-^). Notably, cholesterol^-^/cholesterol^-^ show a recovery rate of about 40 %, the same as *scl-12&13;daf-2* mutants (fig. 3D). Surprisingly, however, we found that *daf-2* recover normally when entry plates are cholesterol^+^ and exit plates are cholesterol^-^ (cholesterol^+^/cholesterol^-^), indicating that environmental cholesterol during pre-dauer development at L1 and L2d larval stages plays an important role for exiting dauer later. This assumption is further consolidated by the fact that animals, which were grown under cholesterol^-^ conditions and exit dauer on cholesterol^+^ plates (cholesterol^-^/cholesterol^+^), experienced impaired recovery, at the same rate as cholesterol^-^/cholesterol^-^ worms. In *scl-12&13;daf-2* mutants, however, this phenomenon could not be observed as all animals showed the same impaired recovery independently of cholesterol conditions, mimicking the effect of lacking dietary cholesterol in pre-dauer animals (fig. 3D). This indicates that cholesterol, SCL-12 and SCL-13 act within the same mechanism of dauer recovery.

**Figure 3:**
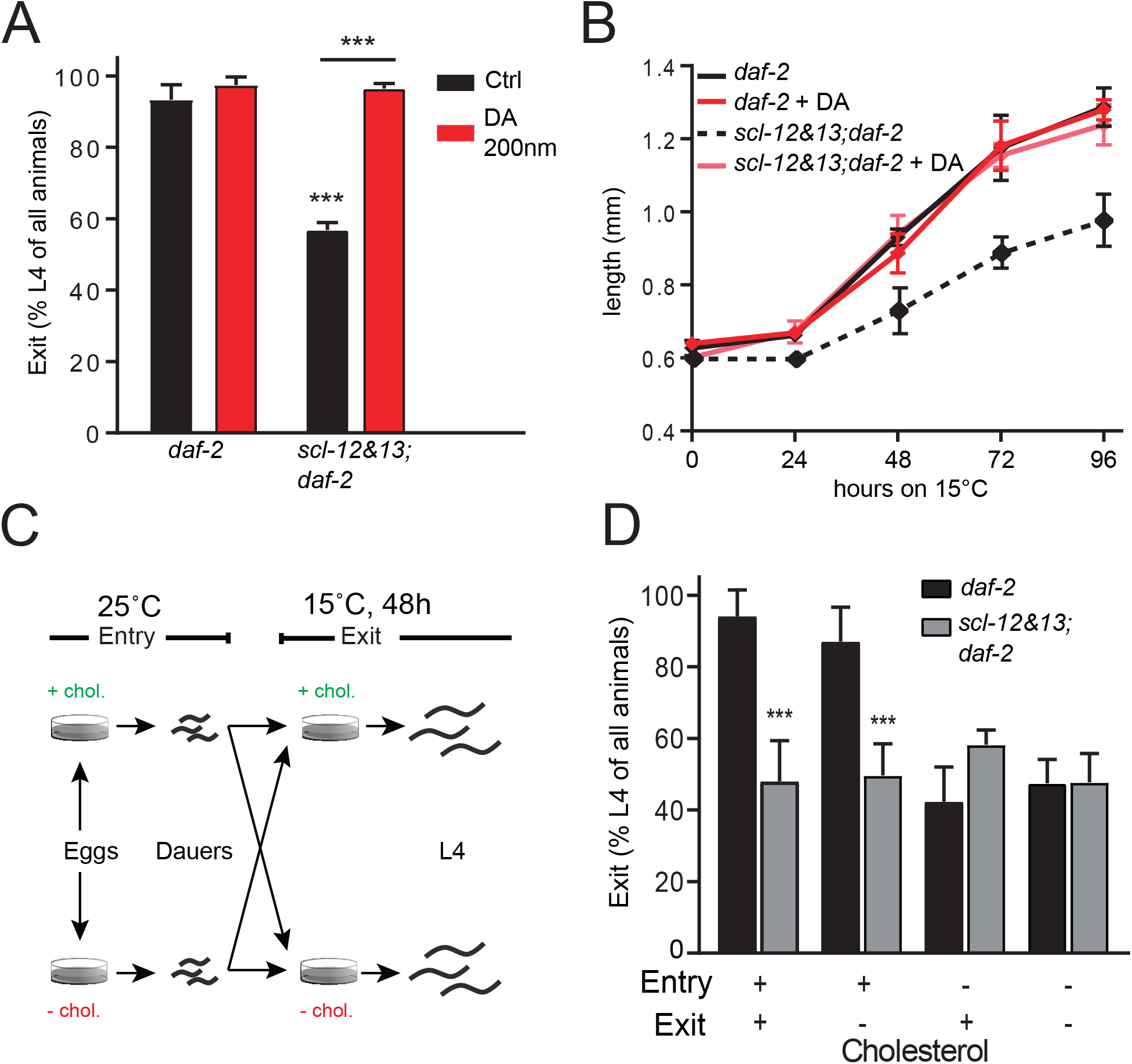
*scl-12&13* mutations lead to cholesterol-dependent impaired dauer exit. **A** Percentage of *daf-2* and *scl-12&13;daf-2* populations that develop into L4 larvae 48 h after dauer recovery was induced by temperature switch to 15°C, control (ethanol; black) and when treated with 200 nm (25S)-Δ7-dafachronic acid (DA; red). **B** Growth in length of *daf-2* and *scl-12&13;daf-2* after induction of dauer exit by temperature switch and the effect of DA thereon. **C** Scheme for experimental procedure in fig. 3D. **D** Percentage of *daf-2* (black) and *scl-12&13;daf-2* (gray) populations that develop into L4 larvae 48 h after dauer recovery was induced by temperature switch to 15°C. Worms were grown with or without cholesterol (+/-) from egg to dauer (“Entry”) and with or without cholesterol from dauer to L4, while exiting dauer (“Exit”).

Dauer exit in *C. elegans* is promoted by dafachronic acids (DAs), which are derivatives of cholesterol. Binding of DAs to DAF-12 switches this nuclear hormone receptor to a dauer-suppressing mode [11]. When supplemented with 200 nM DA, recovery rate and growth during dauer exit at 15°C of *scl-12&13;daf-2* were fully restored (fig. 3A, B). Thus, proper localization and transport of cholesterol in dauer is of importance for biosynthesis of DA during exit.

### Cholesterol is transported from the gut lumen to lysosomes via the endocytic pathway and then further to the epidermal region

As shown above, SCL-12 and cholesterol are moving from the gut lumen to punctate structures during dauer exit, where SCL-12 is degraded. We hypothesized that these structures could be lysosomes. To test this, we co-labeled *daf-2* dauer larvae with TopFluor cholesterol (green) and a Lysotracker dye (red). After 24 h of exit, there is a strong overlap between these markers (fig. 4A). After 48 h, Lysotracker and TopFluor cholesterol do not co-localize anymore and the green fluorescence is seen in concentrated blobs in the epithelial region, which could be lipid droplets. This transport across the pseudocoelomic cavity to the epithelial region could mean that cholesterol undergoes further spatial redistribution and eventually serves for DAF-9-dependent DA synthesis in the hypodermis [30]. Next, we crossed the SCL-12::mScarlet;*daf-2* reporter strain (red) into the well-established genetic lysosomal marker LMP-1::GFP (green) and found a partial overlap of the fluorescent signals during dauer exit (fig. 4B), consistent with SCL-12 and cholesterol moving to the lysosomes during dauer exit.

**Figure 4:**
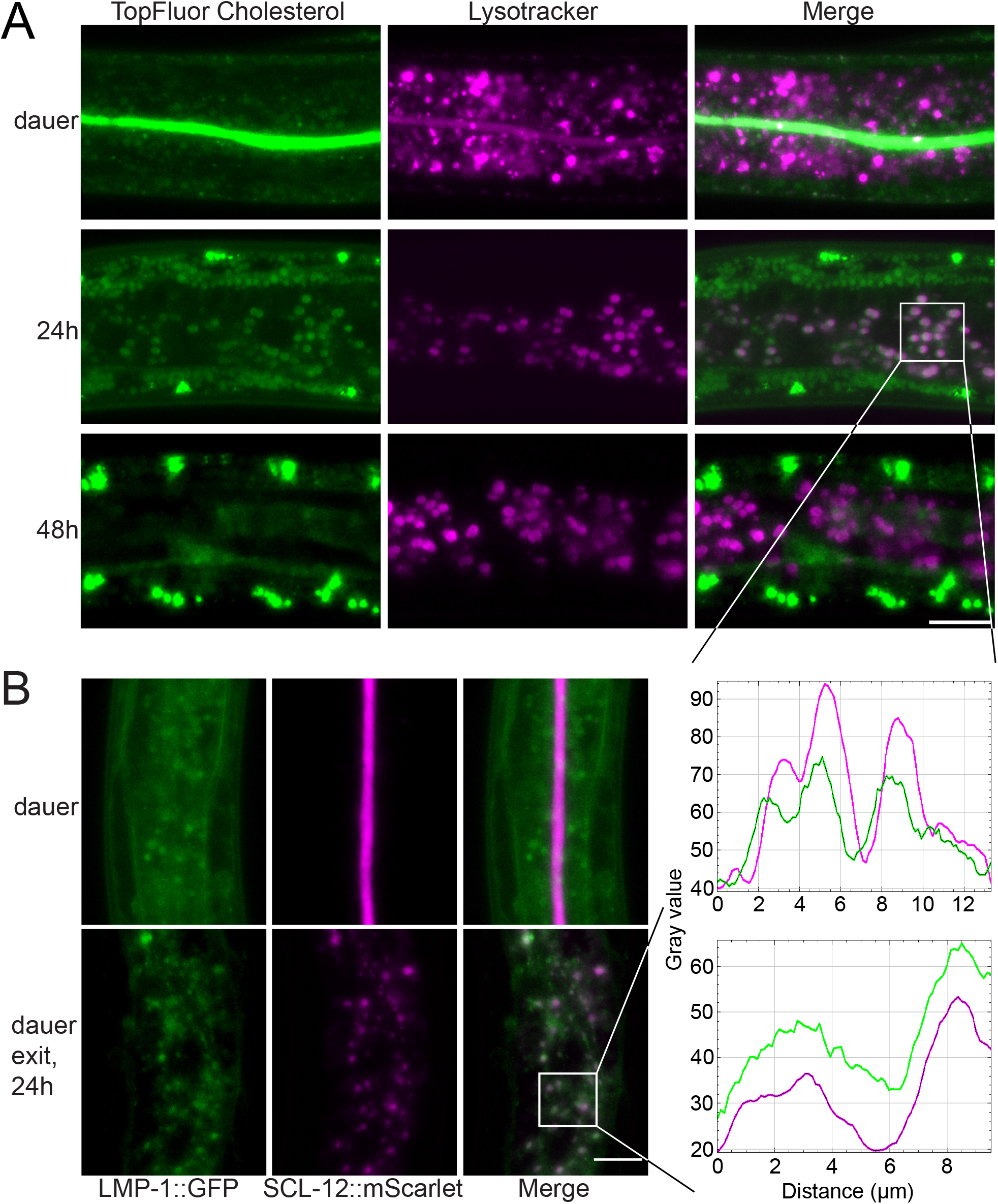
Cholesterol is transported from the gut lumen to intestinal lysosomes with SCL-12 and then further to the epidermal region. **A** *daf-2* dauers co-labelled with TopFluor cholesterol (green) and Lysotracker dye (magenta; top panels), and 24 h and 48 h after induction of dauer exit via temperature switch. Scale bar: 15 μm. The profile (fluorescence intensity in space) of both fluorophores in intestinal cells is similar at 24 h of dauer exit, indicating co-localization. **B** LMP-1::GFP;SCL-12::mScarlet;*daf-2* dauers (top panels) and 24 h after induction of dauer exit (bottom panels). Scale bar: 10 μm. Lysosomal LMP-1::GFP (green) and SCL-12::mScarlet (magenta) show a similar profile at 24 h of dauer exit, indicating co-localization.

Next, we asked which membrane trafficking pathway could be responsible for the transport of SCL-12 and SCL-13 or cholesterol. To identify potential candidates on this end, we measured dauer exit growth in *daf-2* worms fed with RNAis against genes that according to the literature could be involved in vesicle transport and endocytosis: *rme-1, glo-3, vps-32.2, cav-2, rab-5*, and *rab-7* [31–33]. *rab-5* and *vps-32.2* treatment was lethal before worms reached the dauer state and had to be excluded. Whereas *rme-1, cav-2* and *glo-3* had no effect on dauer exit, *rab-7* RNAi showed a strong, highly significant phenotype of impaired dauer exit (fig. S3A). RAB-7 is a late endosomal GTPase responsible for the final lysosomal fusion [34]. The distribution of SCL-12 when worms were fed *rab-7* RNAi indeed showed only a small effect of SCL-12 located within the gut lumen, but a strong effect of SCL-12 enrichment outside of the lumen (fig. S3B). Furthermore, less punctae of SCL-12::mScarlet during dauer exit (12 h) could be identified during *rab-7* RNAi treatment, indicating that SCL-12 is not translocated correctly to the lysosomes (fig. S3B). To visualize the effects on intestinal lysosomes of dauers treated with *rab-7* RNAi we exploited the lysosomal reporter LMP-1::GFP that we crossed with *daf-2* to ease dauer formation. LMP-1 in the mock treatment is being organized in foci, representing lysosomes. Under *rab-7* RNAi treatment, however, LMP-1 seems to be increased in quantity, but diffusely distributed throughout the cytosol or aggregated in larger structures (fig. S3C). Given the strong impairment of dauer exit caused by *rab-7* RNAi and the lack of SCL-12 enrichment in the lysosomes, we propose that the formation of functional lysosomes is an essential process for dauer exit. Since SCL-12 and SCL-13 supposedly deliver cholesterol to the intestinal lysosomes during dauer exit, we wondered whether the lack of functional lysosomes impairs access to sterol supply. Interestingly, when *rab-7* RNAi treated *daf-2* animals were fed a high cholesterol diet, their growth deficit during dauer exit could be partially rescued (fig. S3D), suggesting indeed a crucial role of cholesterol at the lysosomes while exiting the dauer state.

### SCL-12;SCL-13-transported cholesterol activates mTORC1 at the lysosomes to boost growth during dauer exit

We next asked whether the cholesterol that is delivered to the lysosomes by SCL-12 and SCL-13 might activate mTORC1 at the lysosomal membrane, as shown by Castellano et al. *in vitro* [16], during dauer exit to push protein synthesis and growth. Since mutations in the mTORC1 component DAF-15/Raptor or *daf-15* RNAi coordinately suspend development in the L3 stage, we exploited the auxin-inducible degron (AID) system [35], to generate a conditional knock-out of DAF-15 (*daf-15::mNeonGreen::AID*). We did not explore the TOR kinase LET-363 further because it is also part of mTORC2. AID relies on exogenous expression of the plant F-Box protein TIR1, which mediates the depletion of the degron-tagged targets upon exposure to auxin, a plant hormone. Therefore, we expressed *daf-15::mNeonGreen::AID* in *ieSi57(*P*_eft-3_::TIR1::mRuby*) animals to eliminate DAF-15 function in the soma (hereafter referred to as *daf-15::mNG::AID;TIR1*). We then crossed the resulting strain into the *daf-2* background to easily force dauer formation at 25°C. As reported before in non-dauer larval stages, we found that DAF-15 is localized to the lysosomes in dauers (fig. S4A) [19]. To prove the AID principle, we exposed *daf-15::mNG::AID;TIR1* to 400 μM auxin, and found that the fluorescent tag of DAF-15 mostly disappears within one hour (fig. S4B), and worms synchronized as L1 on auxin arrest at the L3 stage, as reported in *daf-15* deletion mutants (fig. S4C) [36]. We next examined dauer exit, and found that in both *daf-15::mNG::AID;TIR1;daf-2* dauers and *daf-15::mNG::AID;TIR1* starvation dauers treated with auxin exit is significantly impaired as indicated by a strong decrease in growth compared to the control (fig. 5A and S4D). In fact, the animals die prematurely (fig. 5B). Interestingly, OCR still strongly increased during dauer exit, demonstrating that DAF-15-dependent growth and mitochondrial activation/metabolic switching toward OxPhos is not correlative (fig. S4E).

**Figure 5:**
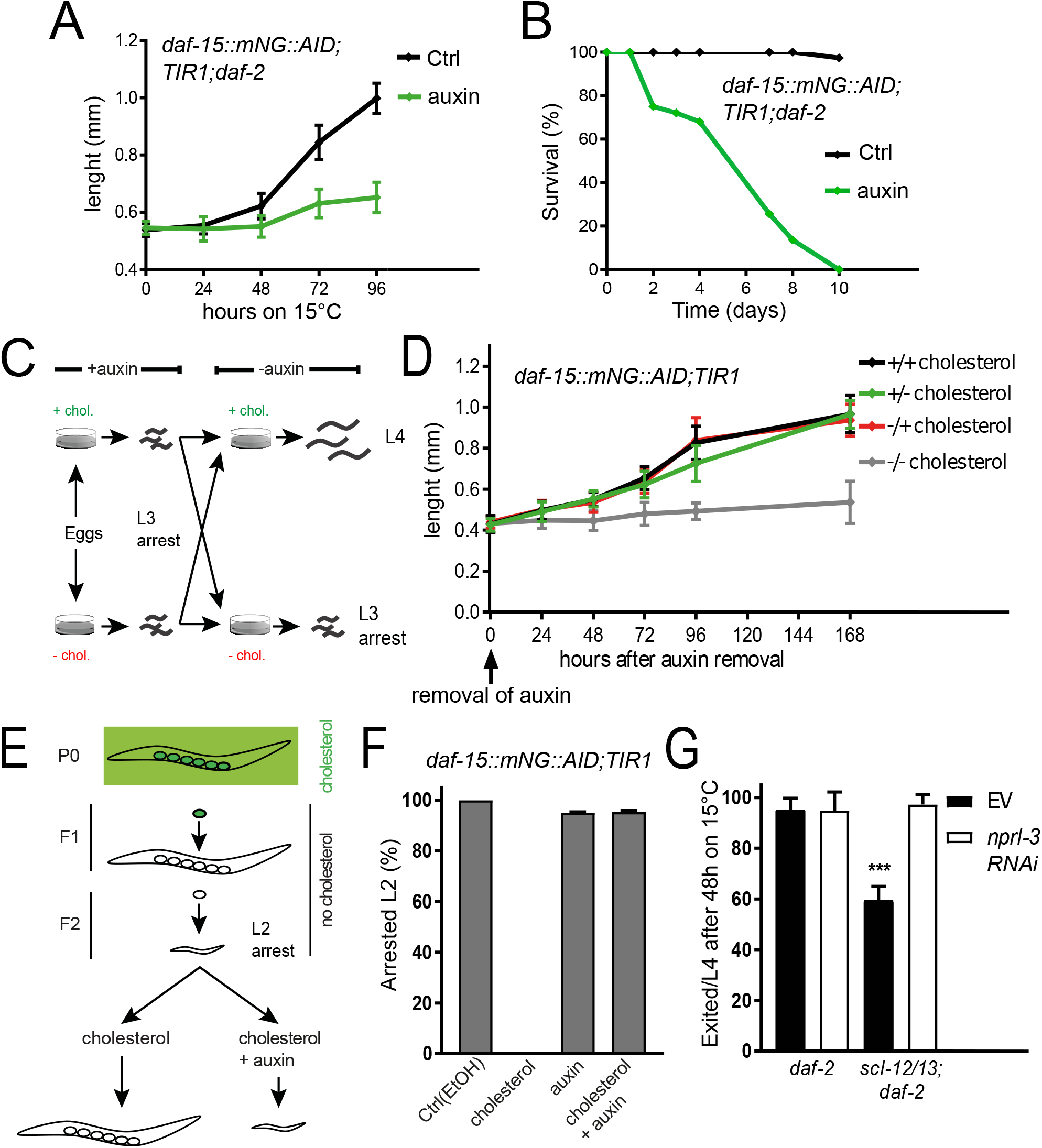
SCL-12;SCL-13-transported cholesterol activates mTORC1 at the lysosomes to boost growth during dauer exit. **A** Growth in length in *daf-15::mNeonGreen::AID;TIR1;daf-2* exiting dauers induced by temperature switch treated with 400 μM auxin (green) or solvent control (ethanol; black). **B** Survival of *daf-15::mNeonGreen::AID;TIR1;daf-2* exiting dauers treated with auxin or solvent control. **C** Scheme of experimental procedure in fig. 5D. **D** *daf-15::mNeonGreen::AID;TIR1* grown on 400 μM auxin and with (+cholesterol) or without cholesterol (-cholesterol) until arrested as L3. Auxin was removed (0 h) and worms started re-growth on +cholesterol or -cholesterol. **E** Scheme of experimental procedure in fig. 5F. **F** *daf-15::mNeonGreen::AID;TIR1* were grown without cholesterol until they arrested in the F2 generation. Graph shows percentage of the population that stays in the arrested state upon addition of cholesterol (0 %), 400 μM auxin, and cholesterol together with 400 μM auxin, compared to the solvent control. **G** Percentage of *daf-2* and *scl-12&13;daf-2* populations that develop into L4 larvae 48 h after dauer recovery was induced by temperature switch to 15°C, mock treatment (EV; black) and on *nprl-3* RNAi (white).

Next, we aimed to investigate if indeed cholesterol and DAF-15/Raptor work together to promote growth *in vivo*. Hence, we exploited the paradigm that *daf-15::mNG::AID;TIR1* arrest at the L3 state, like *daf-15* deletion mutants, when put on 400 μM auxin as L1 larvae. We let worms enter the arrest on either cholesterol^+^ or cholesterol^-^ conditions, as described above (scheme in fig. 5C). After removal of auxin, the animals were allowed to recover on cholesterol^+^ or cholesterol^-^. Interestingly, *daf-15::mNG::AID;TIR1* could exit the auxin-induced arrest and restart growth on all conditions but cholesterol^-^/cholesterol^-^ (fig. 5D), indicating that DAF-15/mTORC1 can only become fully active in the presence of cholesterol. Subsequently, we performed a *vice versa* approach and tested if DAF-15 is required for the rescue of the growth of sterol-depleted worms by addition of exogenous cholesterol. For this, we exploited the paradigm described in [12] and shown in fig. 5E: Worms grown under sterol-depleted conditions will arrest as L2* larvae in the F2 generation, and continue development when cholesterol is restituted to the food. Strikingly, we found that DAF-15 is essential to restart development after addition of cholesterol, because *daf-15::mNG::AID;TIR1* worms on auxin could not exit the arrested L2* stage after the switch to cholesterol^+^ conditions (fig. 5F). To conclude, it seems that under the tested conditions, growth in *C. elegans* is only possible if DAF-15 and cholesterol are both present. Hence, it is very likely that they act within the same cellular process. Data so far indicated that SCL-12/13-mediated cholesterol transport from the gut lumen to the intestinal lysosomes promotes growth partly through activation of mTORC1. To address this hypothesis, we used an RNAi against the gene coding for Nitrogen Permease Regulator Like 3 (*nprl-3*), which has been shown to activate mTORC1 constitutively [37]. The NPRL-2/NPRL-3 complex, similar to its mammalian orthologue GATOR1, represses mTORC1 and loss of either NPRLs causes a robust mTORC1 hyperactivation [38]. If *scl-12&13* mutants partially arrest during dauer exit and this is due to a lack of mTORC1 activation, the downstream hyperactivation of mTORC1 should restore their ability to exit the dauer state. Indeed, 100% of the *scl-12&13;daf-2* animals grown on *nprl-3* RNAi bypassed the partial arrest during dauer exit (fig. 5G). This indicates that SCL-12 and SCL-13 activate mTORC1 during dauer exit by delivering cholesterol to the lysosomes, which is necessary to boost growth after remaining in a quiescence state through the activation of mTORC1 signaling.

## Discussion

In 2010, it was discovered that the lumen of the *C. elegans* dauer gut is entirely filled with a compact multilamellar material, visible in electron microscopy images as concentric circles in cross-sectional cuts [39]. Based on the data in the present study, we hypothesize that this material contains SCL-12, SCL-13, and cholesterol. SCL-12 and SCL-13 are dauer-specific sterol-binding proteins that might sequester cholesterol to store it during the period worms remain in the dauer state. Cholesterol and sterol metabolism are essential for dauer formation and proper recovery from the dauer state to continue progeneration of the population. It seems of advantage to not only rely on the environment to provide exogenous cholesterol but store it in order to have immediate access when exiting dauer. We hypothesize that the sequestration and compartmentalization in the gut lumen is most likely a mechanism to temporally reduce cholesterol signaling. The transcriptional switch utilizing cholesterol that is needed to exit dauer involves the cytochrome P450 DAF-9 and nuclear hormone receptor DAF-12. We show here that the SCL-bound cholesterol most likely associates with DAF-9 as a substrate for DA production, which leads to DA-binding of DAF-12 and a transcriptional switch that specifies reproductive development. The storage of cholesterol during the dauer stage in the gut lumen as opposed to lipid droplets could serve as a means to create distance from the sites of DA synthesis, a process that occurs in the hypodermis. Lipid droplets are also deposited in the hypodermis and a leakage of precursors could result in undesired DA synthesis when environmental cues still suggest the dauer mode.

*C. elegans* dauers are enriched in 4-methylated sterols, constituting about 50% of their total sterol pool. It is therefore probable that these are also ligands of SCL-12 and SCL-13 [13]. DA cannot be produced out of 4-methylated sterols, but they might be active with regard to activate the mTOR pathway, which must be confirmed in the future.

It has furthermore been shown that the reproductive system of *C. elegans* is enriched in cholesterol [40]. There, it might have an additional structural function for the initial germline development upon dauer exit. As the germline develops rapidly during the transition into reproductive mode and many cell divisions take place, worms potentially use the stored cholesterol as building blocks within new cell membranes [41]. Along these lines, the sequestration and absence of cholesterol during dauer might be an effective way to maintain germline quiescence, which is necessary for the reproductive fitness in post-dauer animals [42].

Here, we found evidence for another, not yet described mechanism of cholesterol-driven dauer exit. The activation of mTOR by cholesterol at the lysosomal membrane, as shown before in tissue culture [16] and here for the first time in a living organism, seems also to be involved in effectively exiting dauer. Lately, it has become clear that lysosomes do not only serve as recycling centers, but also as a hub for intracellular distribution of cholesterol and other nutrients. In fact, lysosomes are a sorting station for dietary cholesterol in eukaryotes. In mammalian cells, low-density lipoproteins (LDLs) carrying cholesterol enter the lysosomes via endocytosis and are disassembled in the lumen. From there, a sterol transport system containing the Niemann-Pick C1 (NPC1) and NPC2 proteins that is localized at the lysosome binds free cholesterol and mediates its transport to a variety of cellular compartments, such as the plasma membrane and the endoplasmic reticulum [43]. A variety of signaling processes are initiated at the lysosomal surface, including growth mediated by mTOR. Hormones, growth factors, and nutrients including glucose, amino acids, and a variety of lipids such as cholesterol can activate the mTOR pathway, which occurs in general when environmental conditions are favorable. Subsequently, mTOR regulates transcription factors such as SREBP to drive lipid biosynthesis, induces ribosome biogenesis and protein synthesis, ultimately leading to cellular growth. When conditions are adverse, mTOR signaling is suppressed, resulting in the inhibition of global protein synthesis and significant energy savings [44]. In the light of this paradigm, it makes sense that mTOR signaling is inhibited during dauer and needs an activation signal at the time of exit. In contrast to mammalian cells and similar to yeast, mTORC1 seems to be localized at the lysosomal membrane independently of its activation in *C. elegans* [19, 45, 46], notably also in the dauer larva that is in a quiescent state, as shown here. In mammals, mTORC1 activation by cholesterol requires a lysosomal transmembrane protein, SLC38A9. Additionally, NPC1 binds to SLC38A9 and is capable of inhibiting mTORC1 signaling [16]. Both SLC38A9 and NPC1 have orthologues in *C. elegans*, F13H10.3 and NCR-1/NCR-2, respectively [47, 48]. Like in the mammalian system, NCR-1 and NCR-2 are responsible to direct cholesterol from the lysosomes to other cellular compartments and *ncr-2;ncr-1* knock-out mutants arrest as dauers instead of forming fertile adults as a result of decreased DA production [11, 48]. However, it seems that neither NCR-1/NCR-2 nor F13H10.3 in *C. elegans* are involved in mTOR activation (K.S., T.V.K., unpublished).

Lastly, SCL-12 and SCL-13 are conserved throughout evolution, with orthologues in yeast [24] and human (glioblastoma pathogenesis related 1 (GLIPR2/GAPR1) and cysteine-rich secretory proteins (CRISP2; CRISP3)), and many in between. CRISP2 and CRISP3 are found in the male reproductive tract, especially in sperm. Here, it has been reported that they are involved in sperm maturation and oocyte binding. Notably, cholesterol needs to be removed from the spermatozoa membrane, inducing a variety of processes all needed for a successful capacitation (i.e. the physiological changes needed to penetrate and fertilize an oocyte) [49]. It has also been shown that CRISP2, similar to SCL-12, binds free cholesterol and cholesteryl acetate [24]. It is an intriguing idea that the mechanism of cholesterol sequestration and trafficking described here in *C. elegans* also plays a role in human male fertility. Notably, SCL-12 is not only found in dauers, but also in male *C. elegans* [50]. CRISP3, on the other hand, is dramatically upregulated in prostate cancer, and GLIPR1 is one of the most highly induced transcripts in human gliomas [51]. Whether these proteins also contribute to an activation of growth via mTOR that is involved in malignant processes needs to be further investigated. It is however likely that the processes we describe here may have implications far beyond the mechanisms that govern dauer exit in *C. elegans*.

## Materials and Methods

### Chemicals

All chemicals came from Sigma-Aldrich (Taufkirchen, Germany) unless otherwise specified.

### *C. elegans* maintenance and strains

*C. elegans* were propagated as described before [52]. Briefly, worms were grown on NGM agar plates that were streaked with *Escherichia coli* NA22 as food source at 15°C. The N2 Bristol wildtype strain (wt), *daf-2(e1370)*, the lysosomal reporter LMP-1::GFP*(vkIs2882)*, the TIR1_soma_ expressing strain P_*eft-3*_::TIR1::mRuby (*ieSi57*) and the *E. coli* strain NA22 were provided by the Caenorhabditis Genetics Center (CGC) at the University of Minnesota.

The strains *scl-12&13*, SCL-12::mScarlet and *daf-15::mNeonGreen::AID* were created for this study using the CrisprCas9 system. The compound mutant strains *scl-12&13;daf-2*, SCL-12::mScarlet;*daf-2*, LMP-1::GFP;*daf-2*, LMP-1::GFP;SCL-12::mScarlet, LMP-1::GFP;SCL-12::mScarlet; *daf-2, daf-15::mNeonGreen::AID;TIR1* and *daf-15::mNeonGreen::AID;TIR1;daf-2* were generated by crossing. All genotypes were confirmed by PCR.

Worms were synchronized by hypochlorite treatment of gravid adults [53]. Wt dauers were obtained by SDS treatment of overcrowded, starved worm populations. These populations where washed from the plates and treated with 1% SDS for 30 min, shaking at 30 °C. After that, worms where washed 3x with ddH2O and spread on NGM plates, where surviving dauers separated from dead non-dauers. For *daf-2(e1370)* dauers, eggs where placed on NGM plates and grown at 25 °C for 72 h, in contrast to 15 °C, the restrictive temperature that was used for maintenance of all mentioned strains.

### Growth on sterol-free and high-cholesterol conditions

The growth on sterol-free medium was performed as previously described [12]. Briefly, sterol-depleted medium was obtained by substituting NGM agar with agarose that was extracted three times with chloroform to remove all traces of sterols. NA22 bacteria were grown in sterol-free DMEM medium, pelleted, and resuspended in M9 buffer. For the F2 arrest assay, worms were propagated on these plates for two consecutive generations. Subsequently, synchronized L1 larvae were grown at 25 °C for 72 h in the F2 generation until developmental arrest occurred.

High cholesterol conditions were prepared as described here [14]. Briefly, cholesterol powder was autoclaved and 1 ml E. coli suspension was added to 1 mg of cholesterol crystals. The bacterial/cholesterol suspension was then incubated for 1 h at 37 °C on a rotary shaker at 200 r.p.m. for the cholesterol to solve. NGM-agar plates were then seeded with this bacterial/cholesterol suspension.

### DA treatment

(25S)-Δ7–DA was produced in the laboratory of Prof. H.-J. Knölker and dissolved in ethanol. The stock solution was added to the bacteria to a final concentration of 200 nM, calculated according to the volume of the NGM agar.

### 2-Dimensional difference gel electrophoresis (2D-DIGE)

Wt dauers where obtained by SDS treatment as described above. Dauers were then kept on NGM agar plates without bacteria lawn (Ctrl), or were placed on NGM plates with NA22 lawn and exit via food introduction was induced for 4 h. After that, worms where collected in ddH2O, washed 3 x and snap frozen in liquid nitrogen. Samples where lysed by 5 freezethawing cycles in an ultrasound water bath. Protein content was determined with an RC DC™ Protein Assay (BioRad, Germany). 50 μg of protein per sample were solved in urea lysis buffer and labeled with 250 pmol CyDye DIGE Fluor dyes (GE Healthcare, Germany). Subsequently, 10 nmol L-lysine was used to quench excess dyes and samples were reduced in rehydration buffer (7 M urea, 2 M thiourea, 50 mM DTT, 4% CHAPS). Ampholytes (BioLytes pH 3–10, BioRad, Germany) were added to a total volume of 350 μl. The labeled protein mixture was transferred by passive rehydration for 24 h into an immobilized pH gradient strip (linear pH 3–10). After that, isoelectric focusing for 55–60 kVh in total was performed in a Protean IEF cell (BioRad, Germany) and the gradient strip was equilibrated in equilibration buffer (6 M urea, 50 mM Tris, 130 mM DTT, 2% SDS, 20% glycerol) for 10 min, before being loaded on a 20 cm wide 12% SDS-polyacrylamide gel. The proteins in the strip were separated by SDS-PAGE (200 V, 5 h) and the gel was imaged using a Typhoon 9500 fluorescence imager (GE Healthcare, Germany) at a resolution of 100 μm/pixel for Cy2 at 488 nm excitation, BP 520/40 emission filter, for Cy3 at 532 nm excitation, BP 580/30 emission filter, and for Cy5 at 633 nm excitation, BP 670/30 emission filter. After imaging, Coomassie blue staining was performed and spots of interest were cut out. The proteins in these gel spots were extracted and identified via geLC-MS/MS [54].

### Label-free LC-MS/MS quantification of *C. elegans* proteins

All the reagents used in the experiments are of analytical grade. LC-MS grade solvents were purchased from Fisher Scientific (Waltham, USA); formic acid (FA) from Merck (Darmstadt, Germany), Complete Ultra Protease Inhibitors from Roche (Mannheim, Germany); Trypsin Gold, mass spectrometry grade, from Promega (Madison, USA). Other common chemicals and buffers were from Sigma-Aldrich. The samples for 0 h (dauer) and 24 h (exiting dauers) time points were produced in three biological replicates. Worms were collected from NGM plates with M9 buffer and washed twice, counted, transferred to lysis buffer and snap-frozen in liquid nitrogen. The frozen worms were thawed on ice and crushed using a micro hand mixer (Carl Roth, Germany). The crude extract was centrifuged for 15 min at 13,000 rpm at 4 °C to remove any tissue debris and the clear supernatant was transferred to a fresh Protein Low-Bind tube (Eppendorf, Hamburg, Germany). The total protein content of the samples was estimated using Pierce BCA protein assay kit from Thermo Scientific (Rockford, USA) and 30 μg of total protein content was loaded to a precast 4 to 20% gradient 1-mm thick polyacrylamide mini-gels from Anamed Elektrophorese (Rodau, Germany). Proteins were in-gel digested and analyzed by LC-MS/MS on a hybrid Q Exactive tandem mass spectrometer as described in [55]. Peptide matching was carried out using Mascot v.2.2.04 software (Matrix Science, London, UK) against *C. elegans* proteome downloaded from Uniprot (November 2020), to which common protein contaminants (e.g. human keratins and other skin proteins, bovine trypsin, etc.) have been added. The precursor mass tolerance and the fragment mass tolerance were set to 5 ppm and 0.03 Da, respectively. Fixed modification: carbamidomethyl (C); variable modifications: acetyl (protein N terminus), oxidation (M); cleavage specificity: trypsin, with up to 2 missed cleavages allowed. Peptides having the ions score above 15 were accepted (significance threshold p < 0.05). The label-free quantification and subsequent statistical analysis were performed using MaxQuant (v 1.6.0.16) and Perseus (v1.6.2.1), respectively.

### *In vitro* sterol binding assay

SCL-12 containing an C-terminal polyhistidine tag was expressed in Sf9 insect cells using the *Baculovirus* expression system and purified using standard nickel-based affinity chromatography followed by size-exclusion chromatography. *In vitro* binding was performed in a 96-well micro assay plate with black walls and clear bottom (Greiner Bio-One, Kremsmünster, Austria) in triplicates. The purified protein was solved in PBS at a concentration of 0.2 mg/ml, and 10x fatty acid binding buffer was added according to the individual volume to reach a final concentration of 20 mM Tris (pH 7.5), 30 mM NaCl, and 0.05% Triton X-100. 10 μl of this protein solution was incubated with different concentrations of TopFluor cholesterol (0, 10, 50, 100 nmol in EtOH) for 1 h at room temperature, or pre-incubated with 50 or 100 nm cholesterol for 30 min. Meanwhile, 12 μl Q-Sepharose beads (Cytiva, Marlborough, USA) were prepared to separate the protein from the unbound ligand. Beads were washed 2x with lipid binding buffer, then the sample was added for 1 h at room temperature. Beads with bound protein were washed 3 x with clean binding buffer to remove unbound ligand. After that, fluorescence was measured with a Tecan Spark fluorometer (Männedorf, Switzerland).

### Microscopy

For fluorescence microscopy, worms were placed on glass slides (Superfrost Plus by Thermo Scientific, Waltham, USA) mounted with 1.5 % agarose pads and 5 mM levamisole in M9 buffer was used for immobilisation. Samples were covered with coverslips (0.17 ± 0.005 mm; Menzel Gläser). Fluorescence microscopy was conducted with a Zeiss LSM 880 upright microscope and Zeiss 10x/0.45 Plan-Apochromat/Air and 63x/1.3 LCI Plan-Neofluar, W/Glyc, DIC objectives. Dauer exit, pumping, and growth determination was performed with an Olympus SZX16 stereomicroscope and a QImaging camera.

A minimum of fifteen animals were analyzed in each of at least 3 independent experiments per condition and two-tailed t-test was performed to determine significance.

### Oxygen consumption assay (OCR)

To measure oxygen consumption of *C. elegans*, a Seahorse XFe96 system (Seahorse Bioscience, North Billerica, USA) was used as previously described [56]. *daf-2* and *scl-12&13;daf-2* were grown on NGM plates for 72 h at 25 °C to reach the dauer state. After that, the worms were removed from the plates, washed tree times with M9 buffer and transferred to fresh plates at 15°C to induce dauer exit. After the respective time of exit, worms were collected and washed at least three times to remove all bacteria and debris. About 100 worms/well were pipetted into a 96-well Seahorse XFe assay plate and OCR was measured at a temperature of 25°C. Subsequently, the worms were removed from the wells and BCA protein determination (Thermo Scientific™ Pierce™ BCA Protein Assay Kit) was performed and used for normalization. 8 wells (technical replicates) were used for each strain and condition and 3 independent biological experiments were performed.

### Dauer survival, dauer exit rate, post-dauer fertility and pumping

*daf-2* and *scl-12&13;daf-2* dauer larvae were prepared by growing the worms on solid medium at 25°C as described above. After collection, dauers were washed three times with M9 buffer and subsequently transferred into 15 ml centrifuge tubes (Corning, NY, USA) containing 10ml sterile M9 buffer supplemented with streptomycin (50 μg/ml) and nystatin (10 μg/ml) at a density of 500 worms/ml. The temperature was kept at 25°C and tubes were under constant agitation. Survival rate was determined in percentage by counting alive and dead dauers in 100 μl aliquots every 2 days.

Dauer exit rate and post-dauer fertility determination was performed as follows. *daf-2* and *scl-12&13;daf-2* dauers were removed from plates, washed three times and 1 dauer per well was allowed to exit the dauer state on fresh 12-well plates at 15°C. For dauer exit rate, plates were screened every day for worms reaching the L4 state, which was counted as “exited”. Worms that remained in dauer or died before reaching L4 were considered “non-exited”. Animals that escaped were censored. For post-dauer fertility determination, plates were checked for eggs and alive progeny every day following exit induction. At least 24 worms per strain were monitored within one biological replicate, and due to a large number of escaping dauers, a minimum of 12 were taken into analysis.

Dauer exit was also induced as described above to monitor the start of pumping, where worms were transferred on 6 cm NGM plates. An Olympus SZX16 stereomicroscope was used to screen about 100 animals for pumping movements in their pharynx at 6, 12, 18, and 24 h after dauer exit was induced. The pumping rate of exiting *daf-2* and *scl-12&13;daf-2* dauers was determined in pumps per min 12, 24, and 48 h after induction of dauer exit. A minimum of three biological replicates was performed for each assay.

### TopFluor cholesterol and Lysotracker Red labelling of *C. elegans*

TopFluor cholesterol was resolved in ethanol at a concentration of 6.5 mM, mixed with bacteria solution and spread on top of cholesterol-free NGM plates at a concentration of 6.5 μM, calculated using the volume of the NGM agar. TopFluor cholesterol plates were used instead of standard NGM plates to grow *daf-2, scl-12&13;daf-2*, and SCL-12::mScarlet;*daf-2* at 25 °C until they reached the dauer state. For dauer exit, worms were transferred onto standard NGM plates at 15°C without TopFluor cholesterol to monitor the distribution of the cholesterol that was sequestered during the dauer state.

LysoTracker™ Deep Red dye (Invitrogen™, Carlsbad, USA) was resolved in DMSO at a concentration of 1 mM and mixed with *E. coli* to a final concentration of 1 μM in the bacteria solution. Lysotracker-containing bacteria were spread on standard NGM plates and allowed to dry in the dark. Eggs were put onto those plates and worms were allowed to enter dauer. Plates were kept in the dark to avoid photobleaching. When dauer exit was induced, the animals were transferred onto fresh standard NGM plates without dye. Mean fluorescence intensity of TopFluor cholesterol, LysoTracker™ Deep Red, and SCL-12::mScarlet was measured using the Mean gray value analysis tool in FIJI [57]. Fluorescence profiles were prepared using the FIJI Plot profile tool.

### RNAi/AID experiments

RNAi experiments were performed as described in Kamath et al. [58]. RNAi clones (*E. coli* HT115) of *cav-2*(C56A3.7), *rme-1*(W06H8.1), *glo-3*(F59F5.2), *vps-32.2*(C37C3.3), *rab-5*(F26H9.6), *rab-7*(W03C9.3), and *nprl-3*(F35H10.7) were taken from the ORFeome RNAi library (Open Biosystems) and experimentally compared to an empty vector clone (L4440). Sequencing to confirm the correct clone was performed before usage. RNAi bacteria was spread and allowed to dry on NGM plates containing 1 mM isopropyl-ß-D-thiogalactopyranoside and 50 μg/ml carbenicillin. After a minimum of 24 h, plates were used for experiments.

Auxin treatment was performed as described here [35]. A 400 mM stock solution of the natural occurring auxin indole-3-acetic acid (Alfa Aesar/Thermo Scientific Chemicals, Haverhill, USA) in ethanol was prepared and NGM plates were poured containing 400 μM auxin.

## Figure legends

**Figure S1: Characterization of SCL-12 expression during dauer entry and exit**

**A** Genomic engineering of *scl-12&13* knock-out mutants and SCL-12 reporter animals. **B** SCL-12::mScarlet;*daf-2* reporter animals expressing SCL-12::mScarlet as indicated by appearance of fluorescence after synchronization in % per population. **C** Representative fluorescent micrographs of SCL-12::mScarlet;*daf-2* reporter animals while entering dauer at 44 h, 50 h and 56 h after synchronization of eggs. 44 h and 50 h insets are brightness adjusted images for increased visibility. Scale bar: 15 μm. **D** SCL-12::mScarlet reporter starvation dauers (wt) while exiting dauer at 1.5 h, 3 h and 6 h after introduction of food. 3 h and 6 h insets are brightness adjusted images. Scale bar: 25 μm. **E** Mean fluorescence intensity of SCL-12::mScarlet reporter starvation dauers (0 h) and while exiting dauer via introduction of food. Black: Fluorescence in the gut lumen. Red: Fluorescence outside of the lumen.

**Figure S2: Characterization of *scl-12&13* mutants in dauer state**

**A** Lenght in mm of *daf-2* (black) and *scl-12&13;daf-2* (gray) dauer larvae. **B** Oxygen consumption rate (OCR) in pmol/min/μg protein of *daf-2* and *scl-12&13;daf-2* dauers as worms exit the dauer state. **C** Dauer survival. **D** Time to reach fertility as indicated by egg laying after dauer exit induced by temperature switch to 15°C. **E** Percentage of pumping animals in a population after dauer exit. **F** Pumping rate in pumps/min after dauer exit. Each dot represents an individual worm.

**Figure S3: The RAB-7-dependent endocytic pathway is required for dauer exit**

**A** Growth in length in *daf-2* exiting dauers induced by temperature switch to 15°C on RNAis against factors of the endocytic and lysosome/lysosome-related organelle pathway *cav-2* (green), *rme-1* (blue), *glo-3* (purple) and *rab-7* (orange), compared with mock treatment (EV: empty vector L4440; black). **B** Mean fluorescence intensity of SCL-12::mScarlet;*daf-2* reporter dauers (0 h) and while exiting dauer after temperature switch to 15°C inside (black) and outside (red) of the gut lumen treated with RNAi against *rab-7* (solid line) and EV (dotted line). Representing images of dauer exit 12 h after temperature switch, scale bar: 15 μm. **C** Representing images of dauers of the lysosomal reporter LMP-1::GFP; *daf-2* treated with EV and *rab-7* RNAi. **D** Growth in length in *daf-2* exiting dauers on RNAi against *rab-7* (green) compared to EV (black/gray), treated with 1 mM cholesterol (high cholesterol), compared to standard growing conditions that comprise 13 μM cholesterol.

**Figure S4: Characterization of *daf-15::mNeonGreen::AID;TIR1* animals**

**A** DAF-15::mNeonGreen;*daf-2* reporter dauers (DAF-15::mNG;green) labelled with Lysotracker dye (magenta). The similar profile of both fluorophores indicates co-localization. Scale bar: 5 μm. **B** *daf-15::mNeonGreen::AID;TIR1* dauers treated with 400 μM auxin for 1 h compared to control (ethanol). **C** Growth in length of *daf-15::mNeonGreen::AID;TIR1* worms synchronized as L1 on auxin (400 μM; gray) compared to the solvent control (black). Sizes of developmental stages are indicated in the graph. **D** Growth in length in *daf-15::mNeonGreen::AID;TIR1* exiting starvation dauers treated with auxin (400 μM; green) or solvent control (black). **E** OCR in pmol/min/μg protein in *daf-15::mNeonGreen::AID;TIR1;daf-2* during dauer exit induced by temperature switch, on auxin or solvent control.

## Notes

### Competing Interest Statement

The authors have declared no competing interest.

