## Supplemental figures for "Activation of mTOR by release of extracellular cholesterol stores controls the transition from quiescence to growth in *C. elegans*"

Figure S1 (Schmeisser et al.)

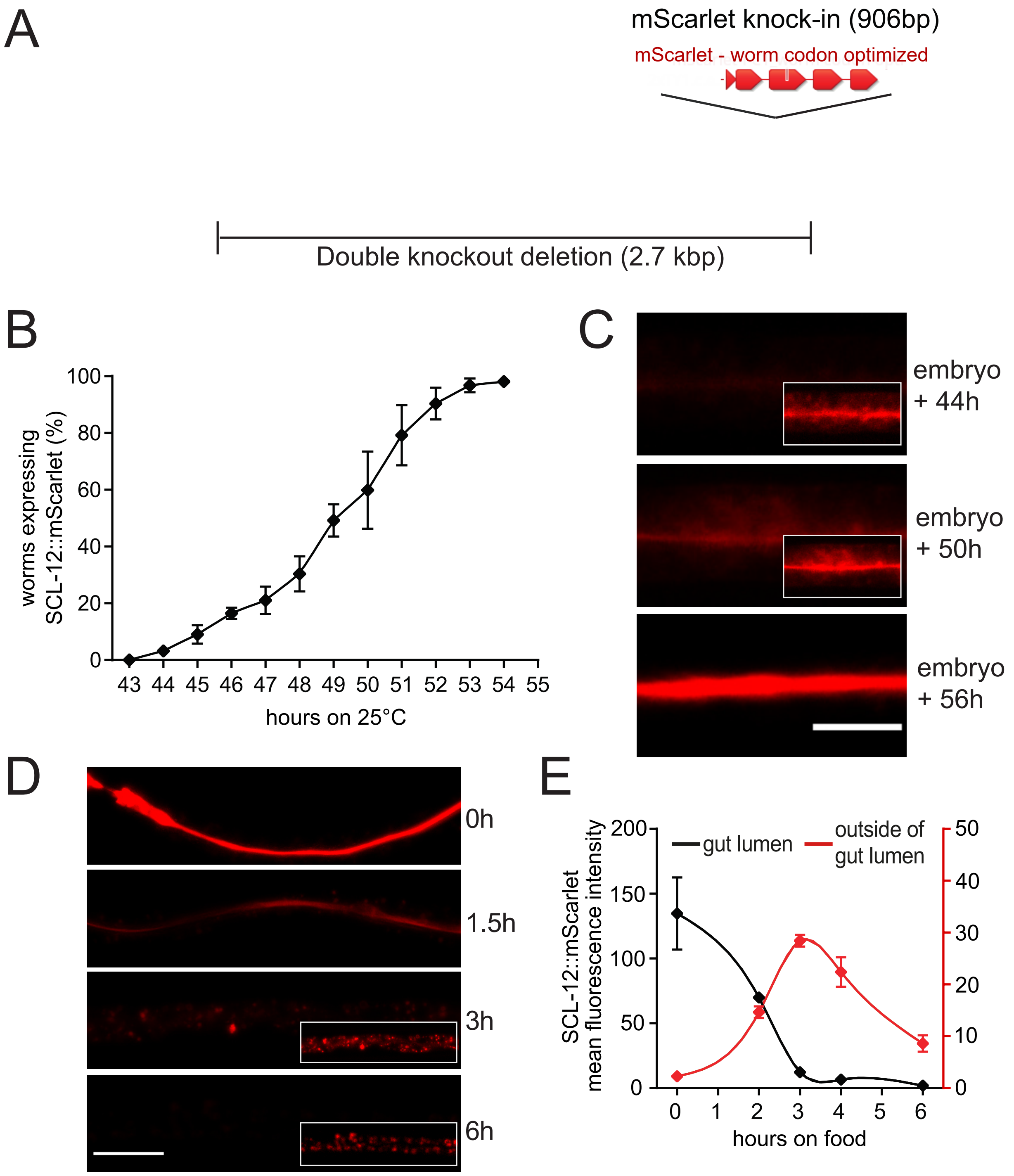

### Figure S2 (Schmeisser et al.)

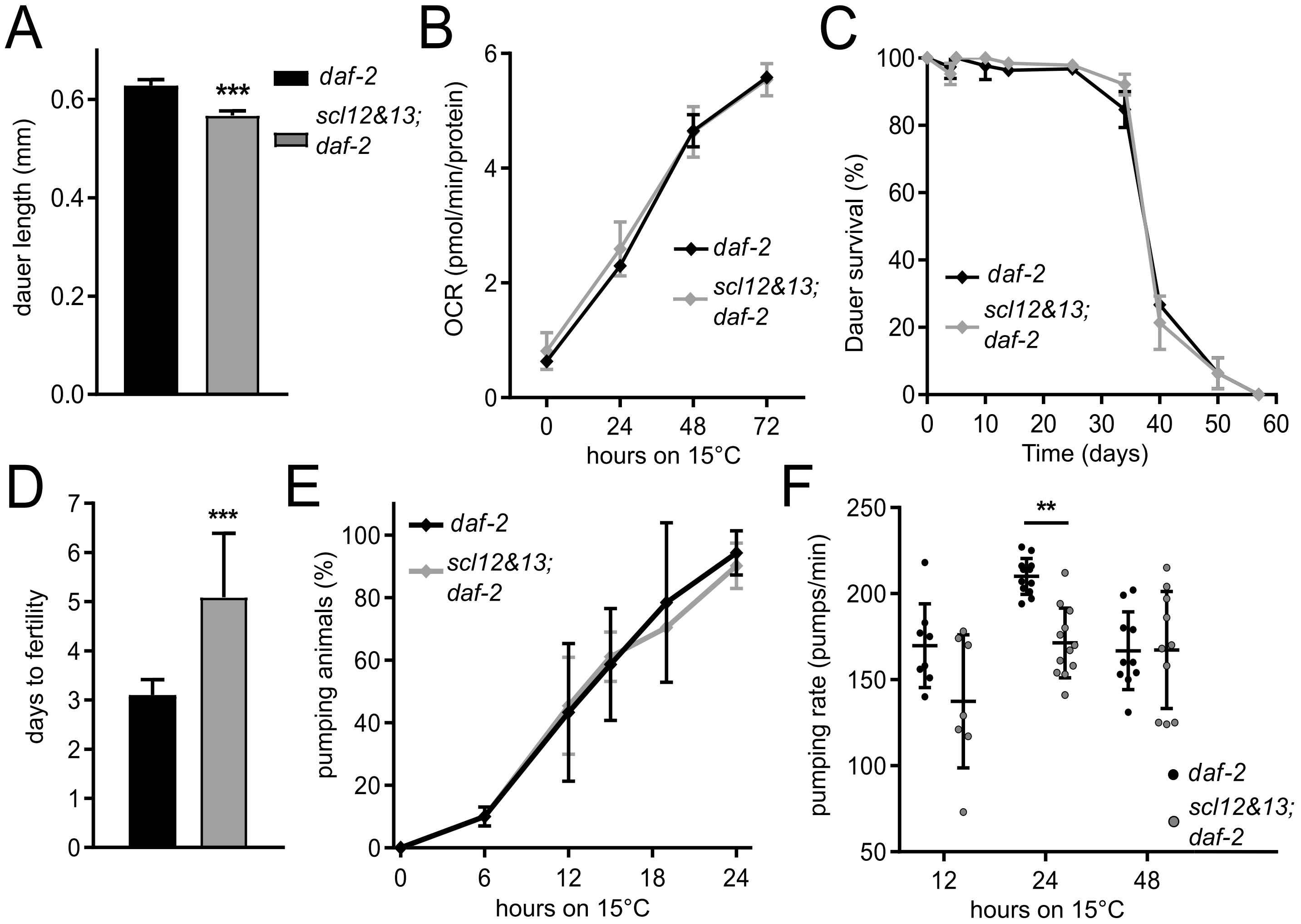

### Figure S3 (Schmeisser et al.)

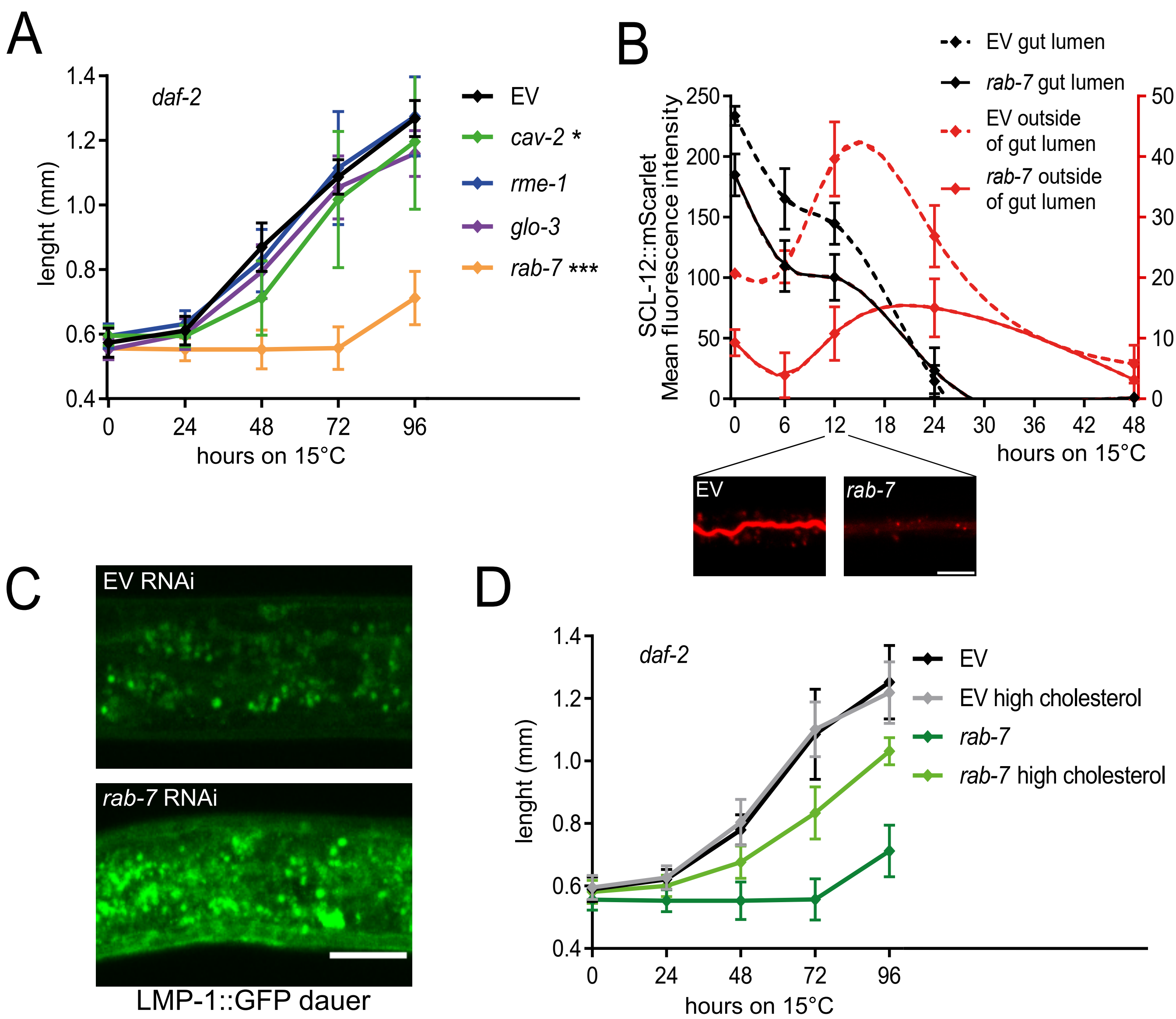

Figure S4 (Schmeisser et al.)

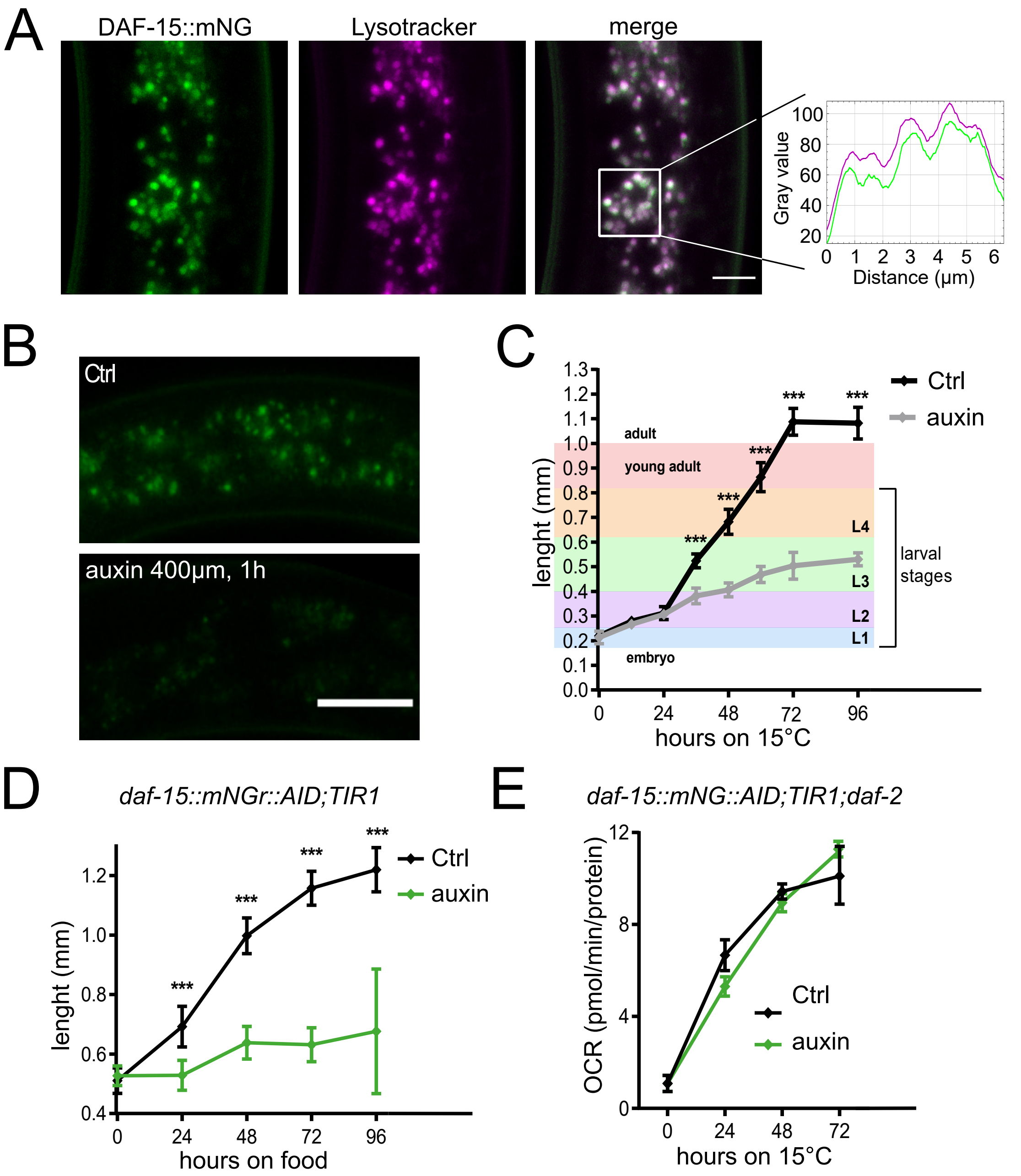
